# Pangenome analysis reveals genetic isolation in *Campylobacter hyointestinalis* subspecies adapted to different mammalian hosts

**DOI:** 10.1101/600403

**Authors:** Daniela Costa, Simon Lévesque, Nitin Kumar, Pablo Fresia, Ignacio Ferrés, Trevor D. Lawley, Gregorio Iraola

## Abstract

*Campylobacter hyointestinalis* is an emerging pathogen currently divided in two subspecies: *C. hyointestinalis* subsp. *lawsonii* which is restricted to pigs, and *C. hyointestinalis* subsp. *hyointestinalis* which can be found in a much wider range of mammalian hosts. Despite *C. hyointestinalis* has been reported as an emerging pathogen, its evolutionary and host-associated diversification patterns are still vastly unexplored. For this reason, we whole-genome sequenced 13 *C. hyointestinalis* subsp. *hyointestinalis* strains and performed a comprehensive comparative analysis including publicly available genomes of *C. hyointestinalis* subsp. *hyointestinalis* and *C. hyointestinalis* subsp. *lawsonii* to gain insight into the genomic variation of these differentially-adapted subspecies. Both subspecies are distinct phylogenetic lineages which present a barrier to homologous recombination, suggesting genetic isolation. This is further supported by accessory gene patterns that recapitulate the core genome phylogeny. Additionally, *C. hyointestinalis* subsp. *hyointestinalis* presents a bigger and more diverse accessory genome, which probably reflects its capacity to colonize different mammalian hosts unlike *C. hyointestinalis* subsp. *lawsonii* that is host-restricted. This greater plasticity in the accessory genome of *C. hyointestinalis* subsp. *hyointestinalis* correlates to a higher incidence of genome-wide recombination events, that may be the underlying mechanism driving its diversification. Concordantly, both subspecies present distinct patterns of gene families involved in genome plasticity and DNA repair like CRISPR-associated proteins and restriction-modification systems. Together, our results provide an overview of the genetic mechanisms shaping the genomes of *C. hyointestinalis* subspecies, contributing to understand the biology of *Campylobacter* species that are increasingly found as emerging pathogens.

## Introduction

The genus *Campylobacter* consists of a diverse group of bacteria currently classified into 29 species and 12 subspecies. Among them, *C. jejuni* and *C. coli* have drawn most of the attention because they are leading causes of human gastroenteritis worldwide^1^. However, the recent application of whole-genome sequencing to study bacterial populations has increased the clinical awareness of campylobacteriosis and highlighted the importance of other neglected *Campylobacter* species, like *C. fetus*^2-4^, as causative agents of human and animal infections. Among them, *C. hyointestinalis* is an emerging pathogen that was first isolated from swine with proliferative enteritis^5^ and has since been sporadically recovered from human infections but also as a commensal from a wide variety of wild, farm and domestic mammals (including cattle, pigs, dogs, hamsters, deer and sheep^6^).

*C. hyointestinalis* is currently divided in two subspecies, namely *C. hyointestinalis* subsp. *lawsonii* and *C. hyointestinalis* subsp. *hyointestinalis*, based on genetic and phenotypic traits^9,10^. While *C. hyointestinalis* subsp. *hyointestinalis* has a broad host range, *C. hyointestinalis* subsp. *lawsonii* is restricted to pigs. Some pioneering studies at both genetic and protein levels have suggested that *C. hyointestinalis* harbors even further intra-species diversity^11-13^ which could facilitate its adaptation to diverse hosts and environments. However, these observations remain to be assessed at higher resolution due to the lack of available genomic data for both subspecies, so the evolutionary forces driving its genetic and ecological distinction have not been explored at the whole-genome level.

Here, we whole-genome sequenced 13 *C. hyointestinalis* subsp. *hyointestinalis* strains isolated from healthy cattle and one strain isolated from a natural watercourse that were sampled on farms located around Sherbrooke, Québec, Canada. By incorporating this information to the available genomes of both subspecies, we performed a pangenome analysis to elucidate the main sources of molecular diversity in both subspecies and the probable genetic mechanisms and functional characteristics that distinguish the host-restricted *C. hyointestinalis* subsp. *lawsonii* from the generalist *C. hyointestinalis* subsp. *hyointestinalis*. Our work provides the first comprehensive analysis of *C. hyointestinalis* subspecies at the pangenome level and will guide future efforts to understand the patterns of host-associated evolution in emerging *Campylobacter* pathogens.

## Results

By whole-genome sequencing 13 *C. hyointestinalis* subsp. *hyointestinalis* strains, we enlarged by 45% the current collection of available genomes for *C. hyointestinalis*. Then, by recovering 29 additional genomes of *C. hyointestinalis* subsp. *hyointestinalis* (n = 19) and *C. hyointestinalis* subsp. *lawsonii* (n = 10) from public databases, we built a genomic dataset consisting of 42 genomes (Table 1). These genomes represent strains isolated between 1985 and 2016 from 5 different hosts in 6 different countries. This dataset was subsequently used to apply comparative pangenomic, phylogenetic and ecological approaches to uncover the main sources of genetic variability in *C. hyointestinalis* subspecies.

**Table 1.** Information of *C. hyointestinalis* genomes analyzed in this work.

| Strain | Subspecies | Country | Date | Host | Material | Reference |
| --- | --- | --- | --- | --- | --- | --- |
| 006A-0059 | Chh | Canada | 2006 | Cow | Feces | This study |
| 006A-0063 | Chh | Canada | 2006 | Cow | Feces | This study |
| 006A-0073 | Chh | Canada | 2006 | Cow | Feces | This study |
| 006A-0091 | Chh | Canada | 2007 | Cow | Feces | This study |
| 006A-0113 | Chh | Canada | 2007 | Cow | Feces | This study |
| 006A-0161 | Chh | Canada | 2007 | Cow | Feces | This study |
| 006A-0170 | Chh | Canada | 2007 | Cow | Feces | This study |
| 006A-0178 | Chh | Canada | 2007 | Cow | Feces | This study |
| 006A-0180 | Chh | Canada | 2007 | Cow | Feces | This study |
| 006A-0191 | Chh | Canada | 2007 | Cow | Feces | This study |
| 006A-0193 | Chh | Canada | 2007 | Cow | Feces | This study |
| 006A-0196 | Chh | Canada | 2007 | Cow | Feces | This study |
| 007A-0283 | Chh | Canada | 2005 | - | Freshwater | This study |
| S1563d | Chh | New Zealand | 2016 | Cow | Feces | Wilkinson et al. (2018) |
| S1564d | Chh | New Zealand | 2016 | Cow | Feces | Wilkinson et al. (2018) |
| S1509d | Chh | New Zealand | 2016 | Cow | Feces | Wilkinson et al. (2018) |
| S1501d | Chh | New Zealand | 2016 | Cow | Feces | Wilkinson et al. (2018) |
| S1597b | Chh | New Zealand | 2016 | Sheep | Feces | Wilkinson et al. (2018) |
| S1603d | Chh | New Zealand | 2016 | Cow | Feces | Wilkinson et al. (2018) |
| S1559c | Chh | New Zealand | 2016 | Cow | Feces | Wilkinson et al. (2018) |
| S1599c | Chh | New Zealand | 2016 | Cow | Feces | Wilkinson et al. (2018) |
| VP28b | Chh | New Zealand | 2010 | Deer | Feces | Wilkinson et al. (2018) |
| VP26b | Chh | New Zealand | 2008 | Deer | Feces | Wilkinson et al. (2018) |
| VP28 | Chh | New Zealand | 2009 | Deer | Feces | Wilkinson et al. (2018) |
| VP30b | Chh | New Zealand | 2011 | Deer | Feces | Wilkinson et al. (2018) |
| S1499c | Chh | New Zealand | 2016 | Cow | Feces | Wilkinson et al. (2018) |
| S1614a | Chh | New Zealand | 2016 | Cow | Feces | Wilkinson et al. (2018) |
| S1592a | Chh | New Zealand | 2016 | Cow | Feces | Wilkinson et al. (2018) |
| S1547c | Chh | New Zealand | 2016 | Sheep | Feces | Wilkinson et al. (2018) |
| LMG-9260 | Chh | Belgium | 1986 | Human | Feces | Miller et al. (2016) |
| DSM 19053 | Chh | United States | 1985 | Pig | Intestine | NCBI |
| ATCC 35217 | Chh | United States | 1985 | Pig | Intestine | JGI |
| RM10074 | Chl | United States | 2009 | Pig | NA | Bian et al. (2018) |
| RM9767 | Chl | United States | 2009 | Pig | NA | Bian et al. (2018) |
| RM9004 | Chl | United States | 2009 | Pig | NA | Bian et al. (2018) |
| RM10071 | Chl | United States | 2009 | Pig | NA | Bian et al. (2018) |
| RM9752 | Chl | United States | 2009 | Pig | NA | Bian et al. (2018) |
| RM9426 | Chl | United States | 2009 | Pig | NA | Bian et al. (2018) |
| RM10075 | Chl | United States | 2009 | Pig | NA | Bian et al. (2018) |
| RM14416 | Chl | NA | 1988 | Cow | Feces | Bian et al. (2018) |
| CHY5 | Chl | United Kingdom | NA | Pig | Stomach | Bian et al. (2018) |
| CCUG-27631 | Chl | Sweden | 1990 | Pig | Stomach | Miller et al. (2016) |

### *C. hyointestinalis* subspecies are genetically isolated lineages

We first reconstructed the species clonal phylogeny starting from a core genome alignment that consisted in 1,320,272 positions (representing 66% of the longest genome), but after removing recombinations only 81,000 positions (representing 6% of the original core genome alignment) remained in the clonal frame. The resulting clonal phylogeny showed a highly structured topology with both subspecies completely separated in two distinct lineages (Fig. 1A, Fig. S1). This observation, together with the clear differences in host distribution suggesting that both subspecies possess isolated ecological niches (Fig. 1B), led us to hypothesize that *C. hyointestinalis* subspecies are undergoing a speciation process driven by host allopatry. Indeed, this was supported by a mean Average Nucleotide Identity (ANI) of ∼95% separating *C. hyointestinalis* subsp. *hyointestinalis* from *C. hyointestinalis* subsp. *lawsonii* (Fig. 1C), which is assumed to be a lower boundary to assign bacterial genomes to the same species^14^. Further evidence supporting the genetic isolation of both subspecies come from exploring genome-wide recombination patterns, which revealed a strong barrier to homologous recombination between *C. hyointestinalis* subsp. *hyointestinalis* from *C. hyointestinalis* subsp. *lawsonii* (with the exception of *C. hyointestinalis* subsp. *hyointestinalis* strains S1499c and 006A-0180 that have recombined with *C. hyointestinalis* subsp. *lawsonii* strains) (Fig. 1D). Furthermore, *C. hyointestinalis* subsp. *hyointestinalis* seems to be much more recombinogenic than *C. hyointestinalis* subsp. *lawsonii,* as evidenced by a significantly higher proportion of their genomes contained within recombinant regions (Fig. 1E). Together, these results indicate that both *C. hyointestinalis* subspecies are separate lineages with considerable genetic isolation probably product of their ecological distinction, as they colonize different mammalian hosts.

**Figure 1.**
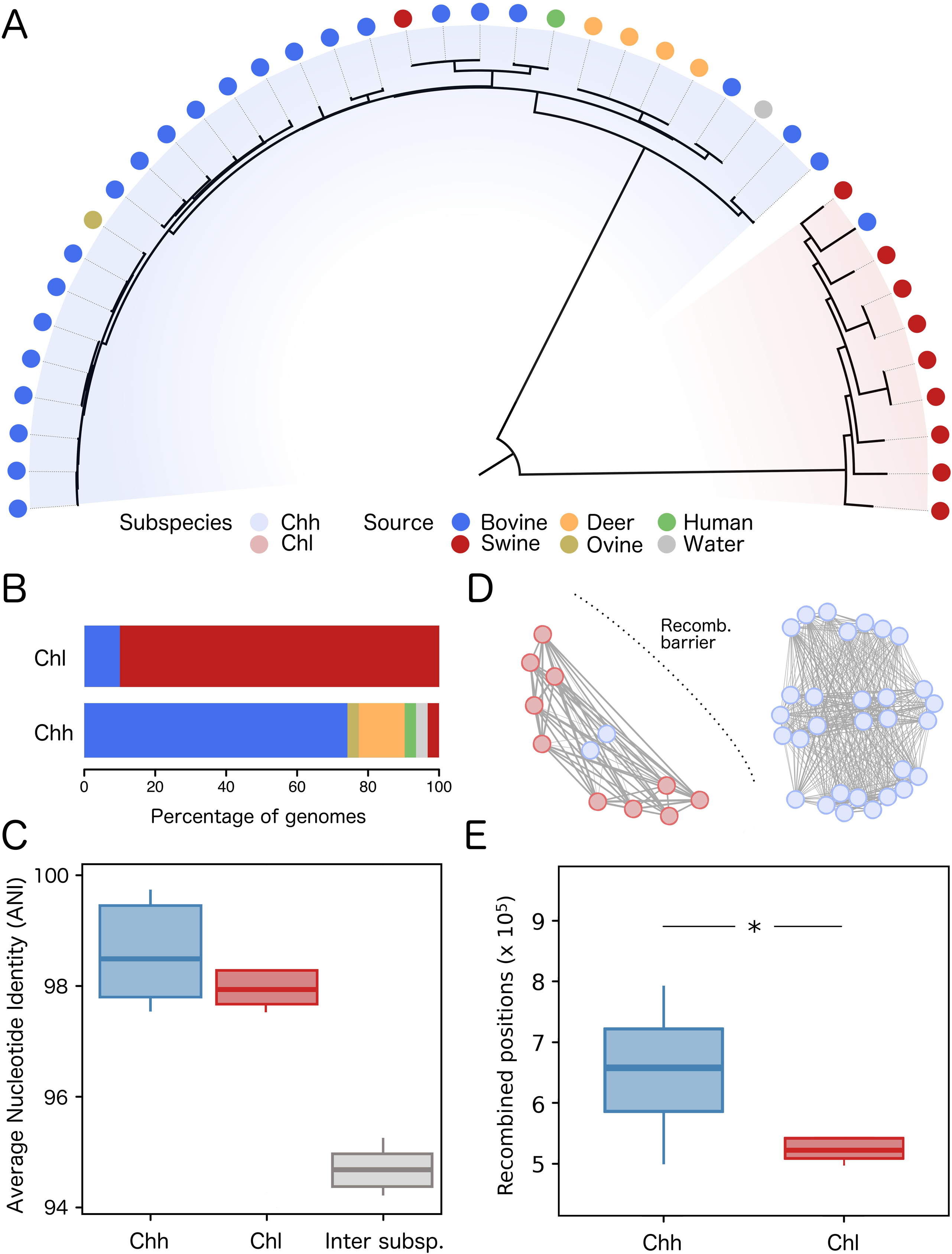
Phylogeny and recombination of ecologically distinct *C. hyointestinalis* subspecies. A) Core genome phylogeny of species *C. hyointestinalis*. Red shade highlights the *C. hyointestinalis* subsp. *lawsonii* lineage and blue shade highlights the *C. hyointestinalis* subsp. *hyointestinalis* lineage. Dots in the tree tips are colored according to isolation source. B) Barplot showing the distribution of hosts in both *C. hyointestinalis* subspecies. C) Boxplots showing ANI values calculated within and between genomes belonging to each subspecies. Inter-subspecies ANI is around 95%, suggesting both subspecies are close to the species definition boundary. D) Network analysis of shared recombinant blocks (edges) between *C. hyointestinalis* genomes (vertexes). A recombination barrier is evidenced between *C. hyointestinalis* subsp. *hyointestinalis* and *C. hyointestinalis* subsp. *lawsonii* given by the lack of recombinant blocks shared between subspecies. E) Boxplots showing the number of recombined positions in the genomes of both subspecies. A statistically significant differences is observed in favor of *C. hyointestinalis* subsp. *hyointestinalis* (p = 0.0035, Mann-Whitney U test).

### Accessory genes discriminate both *C. hyointestinalis* subspecies

To gain further insight on the genomic evolution of *C. hyointestinalis* subspecies we reconstructed its pangenome. A total of 4,317 gene clusters were identified out of which 3,040 (70%) were accessory genes. The accessory genome median size was 580 (IQR = 493-677) and 538 (IQR = 501-575) for *C. hyointestinalis* subsp. *hyointestinalis* and *C. hyointestinalis* subsp. *lawsonii*, respectively. Accordingly, Figure 2A shows a slightly significant difference in the accessory genome size in favor of *C. hyointestinalis* subsp. *hyointestinalis* (p = 0.023, Mann-Whitney U test). This tendency was also observable when calculating the diversity of accessory genes using the inverted Simpson’s index for both subspecies (p = 0.00021, Mann-Whitney U test) (Fig. 2B). Accessory gene presence/absence patterns also allowed to completely discriminate between *C. hyointestinalis* subsp. *hyointestinalis* and *C. hyointestinalis* subsp. *lawsonii* using a Principal Components Analysis, indicating that they have subspecies-specific accessory gene repertories (Fig. 2C). Indeed, 1,562 accessory gene clusters were exclusively found in *C. hyointestinalis* subsp. *hyointestinalis* genomes while only 618 were specific to *C. hyointestinalis* subsp. *lawsonii* genomes. These results support the hypothesis that both subspecies have been diverging isolated from each other for a considerably long time, which probably has impacted the dynamics of their accessory genes and has resulted in specific gene repertories confined to each subspecies.

**Figure 2.**
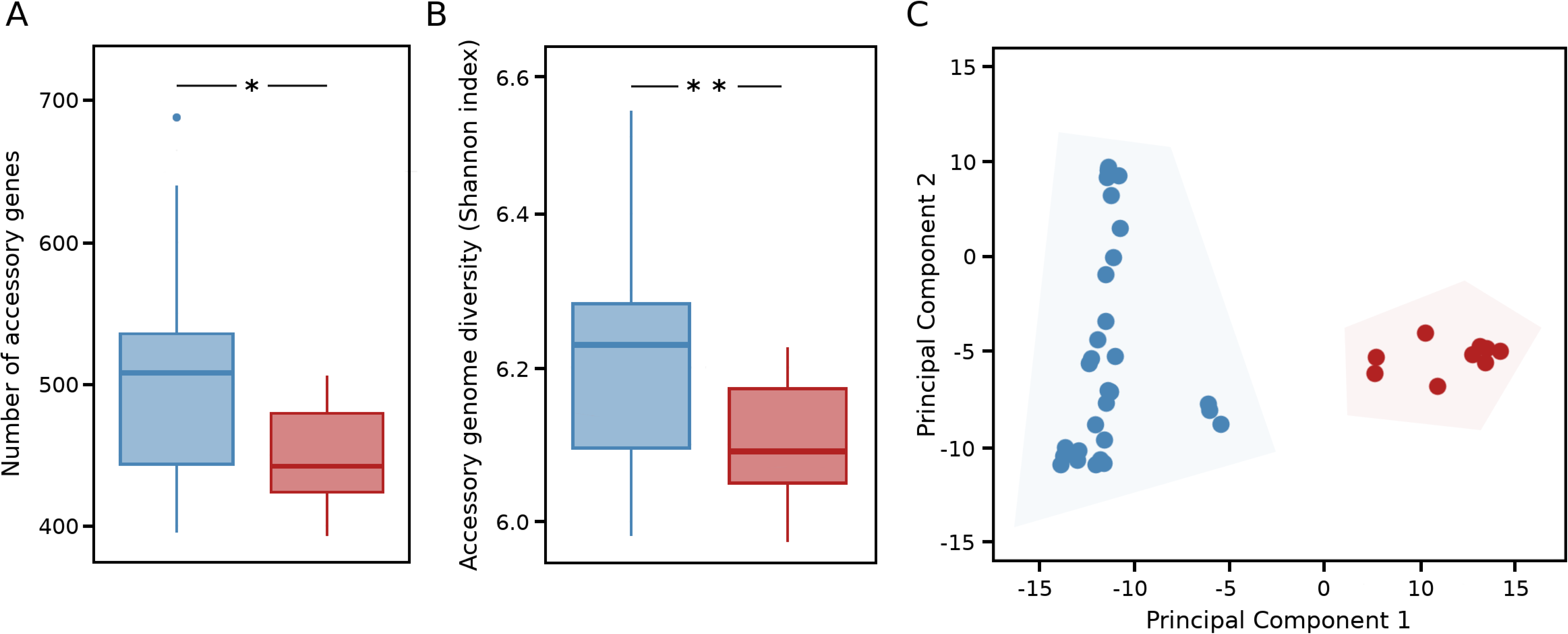
Distinct accessory genomes in *C. hyointestinalis* subspecies. A) Boxplots showing the number of accessory genes (accessory genome size) in both subspecies. *C. hyointestinalis* subsp. *hyointestinalis* possesses a slightly significantly bigger accessory genome than *C. hyointestinalis* subsp. *lawsonii*. (p = 0.023, Mann-Whitney U test). B) Boxplots showing the diversity of accessory genes (as measured by the inverted Simpson index) in both subspecies. *C. hyointestinalis* subsp. *hyointestinalis* has a significantly more diverse accessory genome than *C. hyointestinalis* subsp. *lawsonii*. (p = 0.00021, Mann-Whitney U test). C) Principal component analysis using accessory gene patterns showing that both subspecies represent two completely distinct clusters.

### Functional distinctions in the accessory genome of *C. hyointestinalis* subspecies

To evaluate possible functional aspects associated to the different accessory genomes distinguishing *C. hyointestinalis* subsp. *hyointestinalis* and *C. hyointestinalis* subsp. *lawsonii*, we performed a functional classification of accessory genes based on the eggNOG database^15^. First, we found a complete separation when using functional annotations to perform a Principal Components Analysis (p = 0.001, Permanova test), supporting that accessory genomes are functionally different between both subspecies (Fig. 3A). Then, when looking for those functional categories with greatest contribution to discriminate both subspecies, we found that genes involved in DNA replication, recombination and repair presented the most informative patterns to functionally distinguish *C. hyointestinalis* subsp. *hyointestinalis* from *C. hyointestinalis* subsp. *lawsonii* (Fig. 3B). Given this evidence, we then studied two protein families that are involved in DNA recombination and repair like CRISPR-associated proteins (Cas) and restriction-modification (R-M) systems. Figure 4 shows that both the abundance and diversity of these families in *C. hyointestinalis* subsp. *hyointestinalis* and *C. hyointestinalis* subsp. *lawsonii* present opposite patterns. While R-M systems are significantly more abundant and diverse in *C. hyointestinalis* subsp. *lawsonii* (Fig. 4A-B), Cas proteins are significantly more abundant and diverse in *C. hyointestinalis* subsp. *hyointestinalis* (Fig. 4C-D) (p < 0.01, Mann-Whitney U test). Together, these results indicate that *C. hyointestinalis* subsp. *hyointestinalis* and *C. hyointestinalis* subsp. *lawsonii* genomes harbor distinct molecular machineries involved in DNA recombination and repair, which are probably influencing the differential plasticity observed in their accessory genomes.

**Figure 3.**
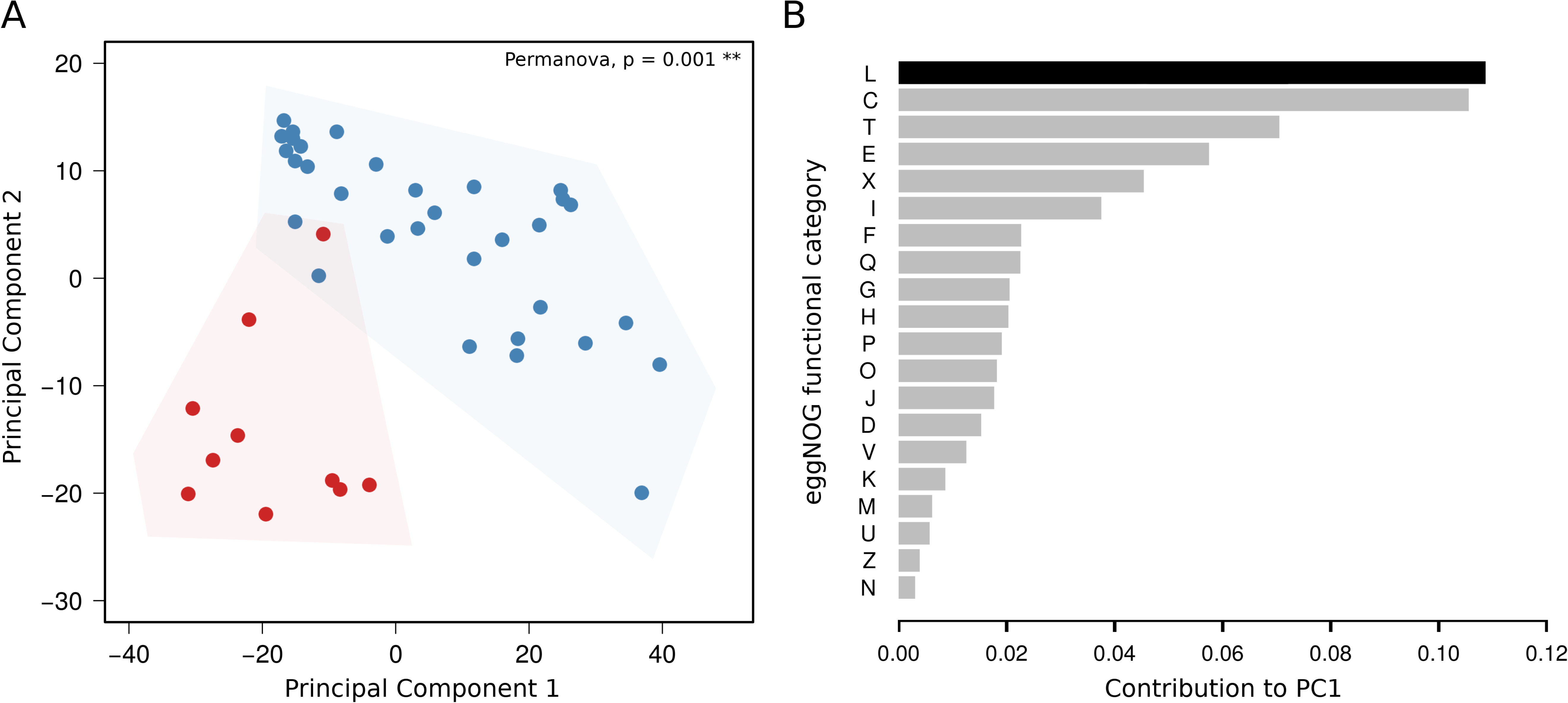
Functionally distinct accessory genomes in *C. hyointestinalis* subspecies. A) Principal component analysis showing that *C. hyointestinalis* subspecies form two different clusters (p = 0.001, Permanova test) based on the functional analysis of their accessory genes. B) Boxplot showing the contribution of each functional category to the variance explained by the first principal component (PC1). Functional category codes resemble those used by the eggNOG database. The top-ranking category (L: recombination and DNA repair) is highlighted in black.

**Figure 4.**
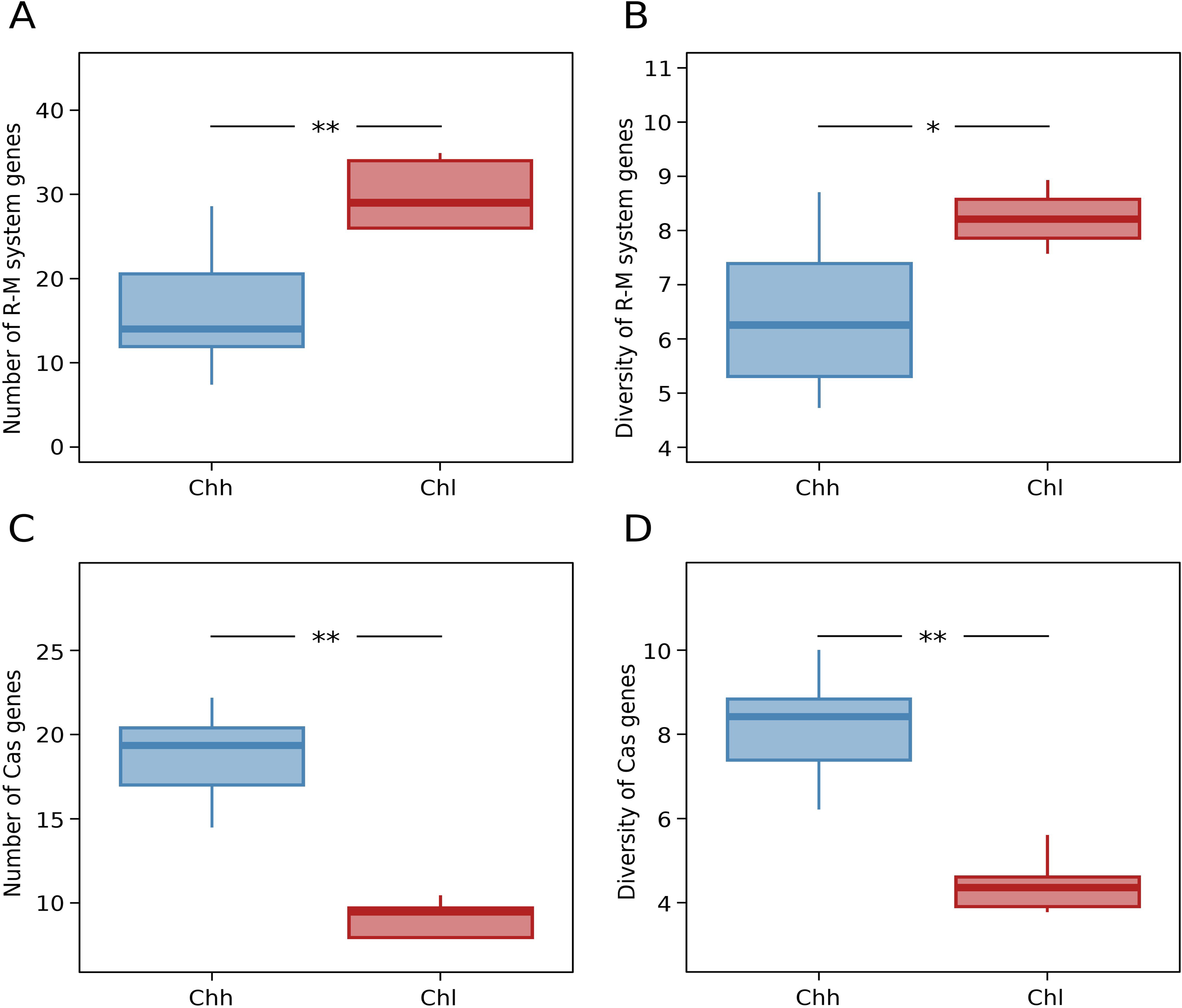
Different repertories of CRISPR/Cas proteins and R-M systems between subspecies. A) Boxplot showing the number of R-M system genes found in *C. hyointestinalis* genomes. A statistically significant difference is appreciated in favor of *C. hyointestinalis* subsp. *lawsonii* (p = 0.00056, Mann-Whitney U test). B) Boxplot showing the diversity (inverted Simpson index) of R-M system genes found in *C. hyointestinalis* genomes. A statistically significant difference is appreciated in favor of *C. hyointestinalis* subsp. *lawsonii* (p = 0.006, Mann-Whitney U test). C) Boxplot showing the number of CRISPR/Cas protein genes found in *C. hyointestinalis* genomes. A statistically significant difference is appreciated in favor of *C. hyointestinalis* subsp. *hyointestinalis* (p = 0.00016, Mann-Whitney U test). D) Boxplot showing the diversity (inverted Simpson index) of CRISPR/Cas protein genes found in *C. hyointestinalis* genomes. A statistically significant difference is appreciated in favor of *C. hyointestinalis* subsp. *hyointestinalis* (p = 0.00021, Mann-Whitney U test). In all cases blue boxes correspond to *C. hyointestinalis* subsp. *hyointestinalis* and red boxes to *C. hyointestinalis* subsp. *lawsonii*.

## Discussion

Recently, the first comparative analysis of multiple *C. hyointestinalis* strains at whole-genome resolution confirmed the previously observed highly diverse nature of this bacterial species. This study revealed a great level of plasticity between *C. hyointestinalis* genomes, with high incidence of recombination and accessory gene gain/loss as the main factors contributing to the observed diversity within this species^16^. However, this study was mainly performed using *C. hyointestinalis* subsp. *hyointestinalis* genomes, including a single representative genome of *C. hyointestinalis* subsp. *lawsonii*. This limitation prevented to compare if the observed trends were conserved between both subspecies or if evolutionary forces are differentially impacting their genomes. Accordingly, our work increased the availability of *C. hyointestinalis* subsp. *hyointestinalis* genomes from a previously unsampled geographic region and by taking advantage of the recent release of novel *C. hyointestinalis* subsp. *lawsonii* genomes, we performed a comparative pangenome analysis that revealed the main forces underpinning the genomic diversity found in *C. hyointestinalis* subsp. *hyointestinalis* and *C. hyointestinalis* subsp. *lawsonii*.

*C. hyointestinalis* subspecies are ecologically distinct since *C. hyointestinalis* subsp. *lawsonii* is restricted to pigs while *C. hyointestinalis* subsp. *hyointestinalis* is a generalist that colonizes several mammalian species. Host specialization has been observed in other *Campylobacter* species, such as in *C. fetus* lineages that preferably infect cows, humans or reptiles^4,17^, in phylogenetically distinct *C. coli* isolates from diseased humans or riparian environments^18^, and in global clonal complexes of *C. jejuni* with differential host preferences^19^. In most of these cases, strong lineage-specific recombination and accessory gene gain/loss patterns have been identified, concordantly to what is expected for bacterial lineages that undergo ecological isolation. For example, the barrier to homologous recombination evidenced between *C. hyointestinalis* subspecies has been also detected between mammal- and reptile-associated *C. fetus* subspecies^17^, and lineage-specific recombination patterns have been found in the *C. jejuni* clonal complex ST-403 that is unable to colonize chicken^20^. Interestingly, this is correlated with the presence of lineage-specific repertories of R-M systems, as well as we observed between *C. hyointestinalis* subspecies. Moreover, other molecular mechanisms involved in genome plasticity like CRISPR/Cas systems are unevenly distributed in agricultural or non-agricultural *C. jejuni*/*coli* genomes^21^, indicating that these systems are differentially present in ecologically distinct niches resembling again the patterns we observed between *C. hyointestinalis* subspecies.

The maintenance of lineage-specific repertories of molecular machineries that modulate genome plasticity is probably an extended mechanism in *Campylobacter*, considering that recombination is an important evolutionary force for the adaptation and acquisition of a host signature in well-known *Campylobacter* pathogens^22^. In general, adaptation occurs in favor of gradual host specialization, but generalism is also widely observed in nature, for example in extremely successful *C. jejuni* lineages that can be found in high prevalence from both agricultural sources or human infections23. A generalist phenotype can be thought as an advantage for bacteria that colonize farm animals, since it allows the subsistence in multiple mammalian species that thieve in close proximity. However, this also represents an increased risk for zoonotic transmission since these animals are usually in contact with humans. Indeed, this scenario is reflected in *C. hyointestinalis* subspecies, given that the generalist *C. hyointestinalis* subsp. *hyointestinalis* has been frequently isolated from human infections in contrast to *C. hyointestinalis* subsp. *lawsonii* that is restricted to pigs and very infrequently reported in humans.

Despite our analysis uncovered the main forces shaping the intra-specific diversity of *C. hyointestinalis* and our results support the observed epidemiological pattern in both subspecies, the integration of a more comprehensive genomic collection from different hosts, geographic regions and clinical conditions must be necessary to deepening our understanding of the genomic evolution in this emerging pathogen and other neglected *Campylobacter* species.

## Methods

### Sampling and bacterial isolation

Samples were collected as described previously. Briefly, cattle feces samples were transported in Enteric Plus medium (Meridian Bioscience Inc, Ohio, USA) and processed on the same day. About 1-2 g of each fecal sample were transferred to 25 ml of Preston selective enrichment broth (Oxoid, Nepean, Ontario, Canada) and incubated 3-4 h at 37° C and then transferred to 42° C and incubated for 48 h. After incubation, 20 μl were streaked on a Karmali plate (Oxoid) and incubated at 42° C for 48 h. For environmental water, 3000 ml of water were collected and transported on ice to the laboratory, held at 4°C and tested within 24 h. Water was filtered through a 0.45 μm pore-size membrane filter and Preston broth and Karmali plate were used as above to isolate *Campylobacter*.

### Whole genome sequencing, available data and taxonogenomic analyses

Cells were pelleted from culture plates and phosphate-buffered saline (PBS). Genomic DNA preparation was performed using a BioRobot M48 (Qiagen). DNA was prepared and sequenced using the Illumina Hi-Seq platform with library fragment sizes of 200-300 bp and a read length of 100 bp at the Wellcome Sanger Institute. Each sequenced genome was *de novo* assembled with Velvet^24^, SSPACE v2.0^25^ and GapFiller v1.1^26^. Resulting contigs were annotated using Prokka^27^. Species membership was checked by calculating the Average Nucleotide Identity (ANI) index as previously described^28^. Available genomic data at the time of designing this work consisted in 19 *C. hyointestinalis* subsp. *hyointestinalis* strains and 10 *C. hyointestinalis* subsp. *lawsonii* strains, that were added to the 13 *C. hyointestinalis* subsp. *hyointestinalis* sequenced in this work to build a final dataset of 42 genomes (Table 1).

### Pangenome and recombination analyses

A multiple genome alignment was performed with the progressiveMauve algorithm^29^ and the final core genome alignment was defined by concatenating locally collinear blocks (LCBs) longer than 500 bp present in every genome. Recombinant regions were identified running Gubbins^30^ with default parameters. The pan-genome was reconstructed using a previously implemented^31^ in-house pipeline (https://github.com/iferres/pewit). Briefly, for every genome, each annotated gene is scanned against the Pfam database^32^ using HMMER3 v3.1b2 hmmsearch^33^ and its domain architecture is recorded (presence and order). A primary set of orthologous clusters is generated by grouping genes sharing exactly the same domain architecture. Then, remaining genes without hits against the Pfam database are compared to each other at protein level using HMMER3 v3.1b2 phmmer and clustered using the MCL algorithm^34^. These coarse clusters are then splitted using a tree-prunning algorithm which allows to discriminate between orthologous and paralogous genes. Standard ecological distances over accessory gene patterns were calculated with the vegan package^35^.

### Analysis of specific gene families and functional categories

Several specific gene families of interest were recovered and analyzed form *C. hyointestinalis* genomes. CRISPR-associated protein (CAS) genes were recovered by running HMMER3 v3.1b2 hmmsearch^33^ against Hidden Markov Models for every single CAS gene type from the CRISPRCasFinder database^36^. The REBASE database^37^ was used to retrieve R-M system genes that were compared to the *C. hyointestinalis* genomes using Blast+ blastp^38^ with an identity >70% and query coverage >70% as inclusion thresholds. Alpha diversity for each gene family in each genome was calculated using the Shannon index as implemented in the vegan package^35^. Functional categories were assigned to annotated genes using the eggNOG database^15^ and the eggNOG-mapper tool^36^.

## Acknowledgments

We acknowledge the Pathogen Informatics and Sequencing groups at the Wellcome Trust Sanger Institute for technical support. We also thank to Mark Stares and Hilary Browne at the Host-Microbiota Interactions Laboratory, Wellcome Trust Sanger Institute, for their technical support. G.I. and D.C. are supported by the Agencia Nacional de Investigación e Innovación (ANII, Uruguay) grant FSSA_X_2014_1_105252. This work received partial financial support from Fondo de Convergencia Estructural del Mercosur (FOCEM) grant COF 03/11, the Wellcome Trust grant number 098051 and the Medical Research Council UK grant number PF451.

## Author Contributions

G.I. conceived the idea and designed the experiments. G.I., D.C., I.F., P.F., S.L. and N.K. performed the experiments and analyzed the data. S.L. collected and provided samples and T.D.L. contributed to data analysis and interpretation. G.I. and D.C. wrote the paper with suggestions from all authors. All authors approved the manuscript prior to submission.

